# Molecular adaptation to folivory and the conservation implications for Madagascar’s lemurs

**DOI:** 10.1101/2021.07.06.451309

**Authors:** Elaine E. Guevara, Lydia K. Greene, Marina B. Blanco, Casey Farmer, Jeannin Ranaivonasy, Joelisoa Ratsirarson, Karine L. Mahefarisoa, Tsiky Rajaonarivelo, Hajanirina H. Rakotondrainibe, Randall E. Junge, Cathy V. Williams, Elodi Rambeloson, Hoby A. Rasoanaivo, Vololonirina Rahalinarivo, Laza H. Andrianandrianina, Jonathan B. Clayton, Ryan S. Rothman, Richard R. Lawler, Brenda J. Bradley, Anne D. Yoder

## Abstract

Folivory evolved independently at least three times over the last 40 million years among Madagascar’s lemurs. Many extant lemuriform folivores exist in sympatry in Madagascar’s remaining forests. These species avoid feeding competition by adopting different dietary strategies within folivory, reflected in behavioral, morphological, and microbiota diversity across species. These conditions make lemurs an ideal study system for understanding adaptation to leaf-eating. Most folivorous lemurs are also highly endangered. The significance of folivory for conservation outlook is complex. Though generalist folivores may be relatively well equipped to survive habitat disturbance, specialist folivores occupying narrow dietary niches may be less resilient. Characterizing the genetic bases of adaptation to folivory across species and lineages can provide insights into their differential physiology and potential to resist habitat change. We recently reported accelerated genetic change in *RNASE1*, a gene encoding an enzyme (RNase 1) involved in molecular adaptation in mammalian folivores, including various monkeys and sifakas (genus *Propithecus*; family Indriidae). Here, we sought to assess whether other lemurs, including phylogenetically and ecologically diverse folivores, might show parallel adaptive change in *RNASE1* that could underlie a capacity for efficient folivory. We characterized *RNASE1* in 21 lemur species representing all five families and members of the three extant folivorous lineages: 1) bamboo lemurs (family Lemuridae), 2) sportive lemurs (family Lepilemuridae), and 3) indriids (family Indriidae). We found pervasive sequence change in *RNASE1* across all indriids, a d_N_/d_S_ value > 3 in this clade, and evidence for shared change in isoelectric point, indicating altered enzymatic function. Sportive and bamboo lemurs, in contrast, showed more modest sequence change. The greater change in indriids may reflect a shared strategy emphasizing complex gut morphology and microbiota to facilitate folivory. This case study illustrates how genetic analysis may reveal differences in functional traits that could influence species’ ecology and, in turn, their resilience to habitat change. Moreover, our results support the contention that not all primate folivores are built the same and highlight the need to avoid generalizations about dietary guild in considering conservation outlook, particularly in lemurs where such diversity in folivory has probably led to extensive specialization via niche partitioning.

## INTRODUCTION

The primates of Madagascar include a remarkable number of folivores (1). The island is home to more than 50 extant species of folivorous lemur – and was home to several additional, now-extinct folivorous species – that are phylogenetically and ecologically diverse, as well as endangered (1,2). Extant species include members of six genera in three families (Figure 1A) that independently converged on folivory at various evolutionary time-depths (3). Nutritional reliance on leaves, which contain plant structural carbohydrates and chemical defenses, is a challenging nutritional strategy (4,5) and requires numerous anatomical, physiological, and behavioral adaptations (6).

**Figure 1.**
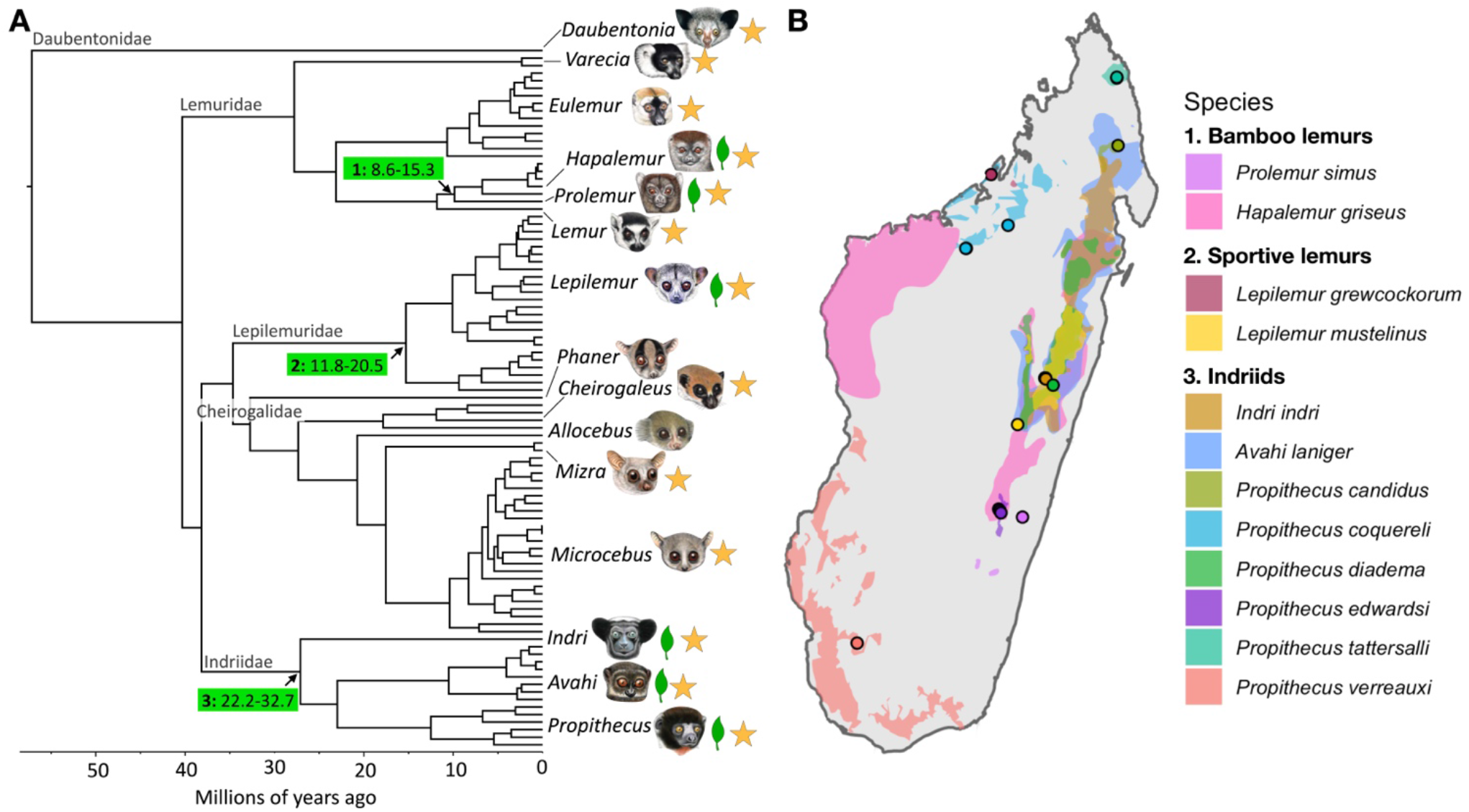
Species included in this study. A = Lemur phylogeny adapted from (3). Illustrations copyright 2020 Stephen D. Nash / IUCN SSC Primate Specialist Group. Used with permission. Arrows denote branches along which folivory emerged, with numbers in the green boxes reflecting the three folivorous lemur groups: 1: bamboo lemurs, 2: sportive lemurs and 3: indriids and number ranges indicating 95% confidence intervals for the time (in millions of years ago) of the basal split in that group from (3). Leaf icons at tips denote folivorous genera and stars denote genera represented in this study. B = Range maps of folivorous species included in this study from IUCN Red List of Threatened Species. Points indicate sampling localities for wild individuals sequenced.

Dietary strategy is of great importance for conservation outlook. Folivores are traditionally considered more resilient to habitat disturbance and change due to the relative abundance of leaves (7–9). However, decades of research indicate that not all folivores are ecologically or physiologically equivalent (10–12). Phylogenetic constraints, for example on body size, the nature of available foliage in a given habitat, and niche partitioning influence which behavioral, anatomical, and physiological routes to folivory different taxa adopt (13,14).

For example, the many folivorous lemurs that live in sympatry (Figure 1B) across Madagascar’s remnant forested ecosystems avoid feeding competition through differentiation in activity patterns, microhabitat use, and selection of leaves with differing amounts of protein, structural carbohydrates, and plant secondary compounds, including tannins and alkaloids (13,15–17). These forms of specialization likely involve varying underlying physiological adaptations to different aspects of leaf digestion, which may in turn influence species’ ability to cope with environmental limitations and change (11,18).

Genetic analysis offers an opportunity to get “under the hood” of different primate folivores to better understand their specific physiological adaptations that facilitate folivory. A prime example is the *RNASE1* gene, which encodes the pancreatic ribonuclease enzyme (RNase1 or RNase A). It shows evidence of adaptive convergence in folivorous mammalian lineages at various evolutionary time depths. It has undergone independent gene duplications followed by rapid sequence evolution in bovine ruminants and colobine monkeys (19–21), who share a common ancestor 85-105 million years ago (Ma) (3,22). Moreover, there have been independent duplications of *RNASE1* followed by parallel amino acid substitutions in both the African and Asian colobine monkey clades, who share an ancestor ~15 Ma (3,20,21). Functionally parallel amino acid substitutions in the duplicated RNase1 peptide in both ruminants and colobines result in a lowered optimal pH for enzymatic activity relative to the conserved peptide (20,21). Although *RNASE1* is expressed across several tissues, it is much more highly expressed in the pancreas of ruminants than that of non-herbivorous mammals (23,24). The duplicated RNase1 is thought to play a key role in ruminant digestion by breaking down RNA produced by the large microbial communities that facilitate cellulose digestion to recover nitrogen for host absorption (21,23). The enzyme’s lowered optimal pH presumably boosts efficiency in the small intestine, which in ruminants has a lowered pH (20,21). Recent work in platyrrhine primates has uncovered evidence for an *RNASE1* duplication in howler monkeys (*Alouatta* spp.) followed by amino acid substitutions and a lowered isoelectric point paralleling those in colobines (25). Isoelectric point (pI) is the pH at which the peptide has no charge, and changes in pI reflect substitutions that alter charge and thus how the enzyme interacts with negatively charged RNA molecules. Howler monkeys are also highly folivorous, but do not have a ruminant digestive system, relying instead on hindgut fermentation (26). The finding of parallel evolution in howler monkeys and colobines thus raises the possibility of an alternative or additional role for RNase1 in hindgut-based foliage digestion.

Our previous work has uncovered positively selected sites in *RNASE1* in sifaka lemurs (genus *Propithecus*) (Guevara et al., 2021). Sifakas, along with the indri (genus *Indri*) and woolly lemurs (genus *Avahi*), belong to the family Indriidae, all of whose members eat diets high in leafy material and show traits associated with obtaining nutrients from leaves. Such traits include shearing teeth and an enlarged caecum and complex colon to facilitate hindgut fermentation using commensurate specialized microbiota (27–30).

In this study, we expanded our analysis to include additional folivorous lemurs to determine if they also show convergent changes in the *RNASE1* gene. Specifically, we analyzed *RNASE1* sequence data for 21 species of lemur, representing all five lemur families (Table 1), many of which are sympatric in eastern Madagascar (Figure 1B).

**Table 1.**
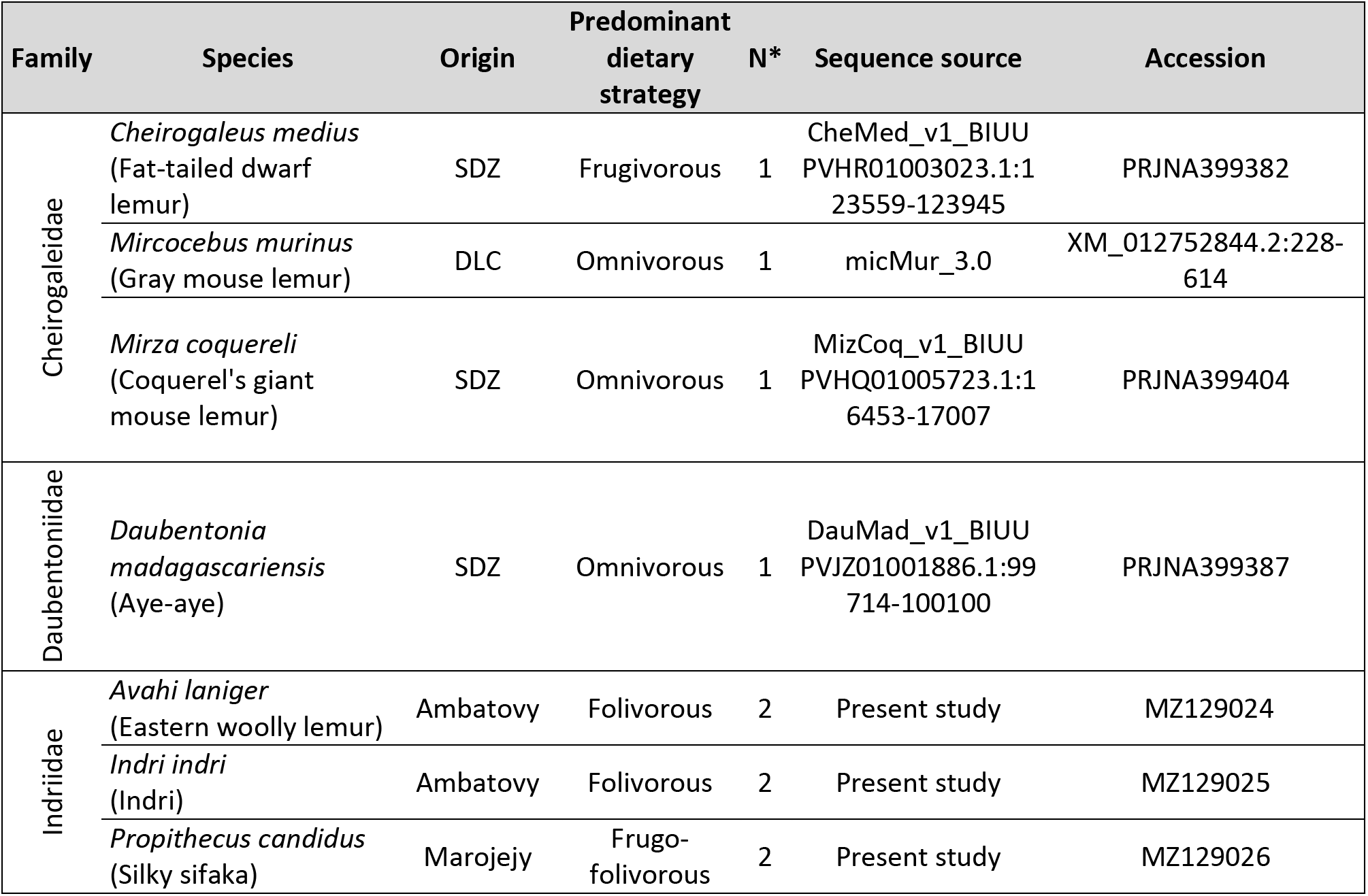
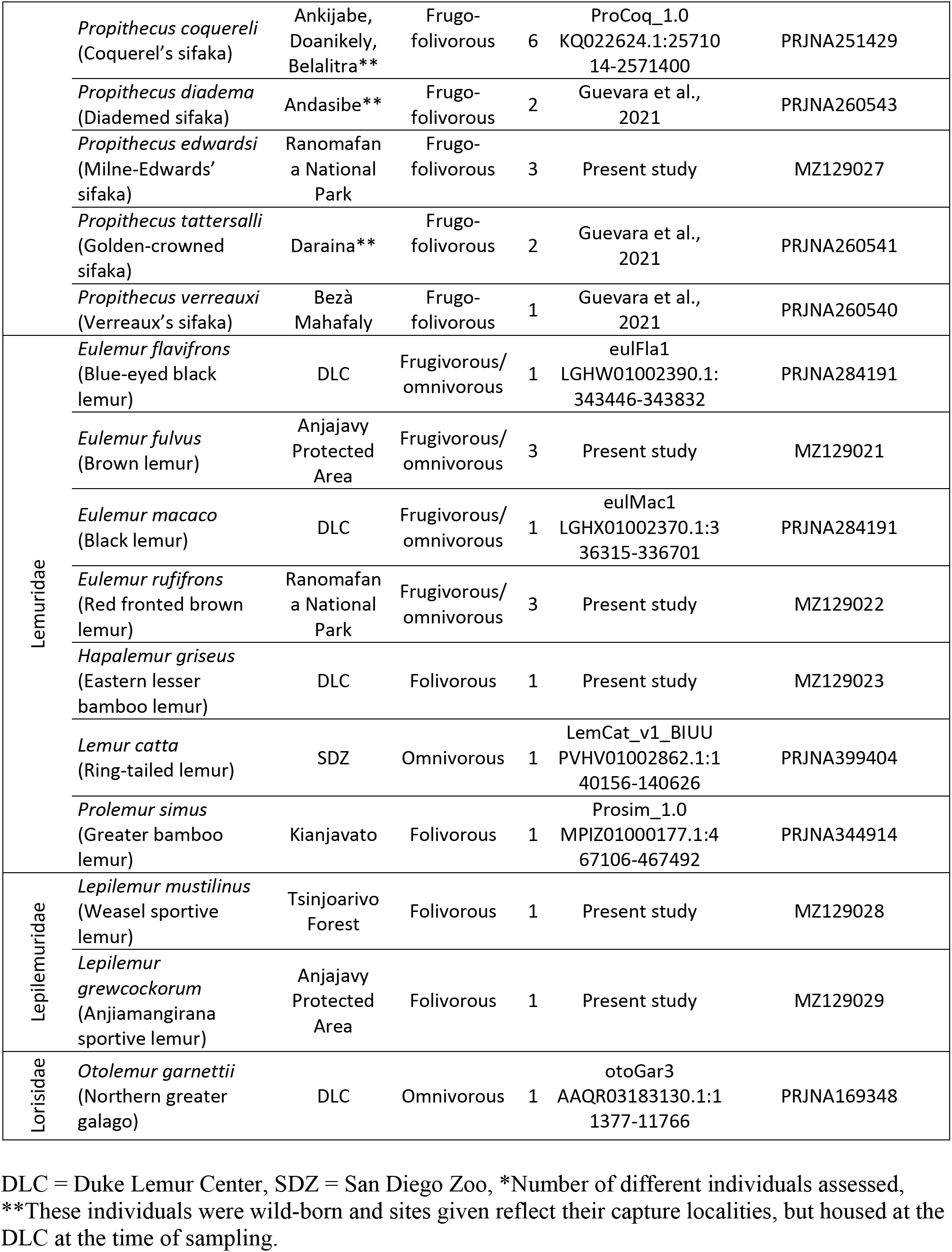
Species and samples included in this study.

**Table 2.**
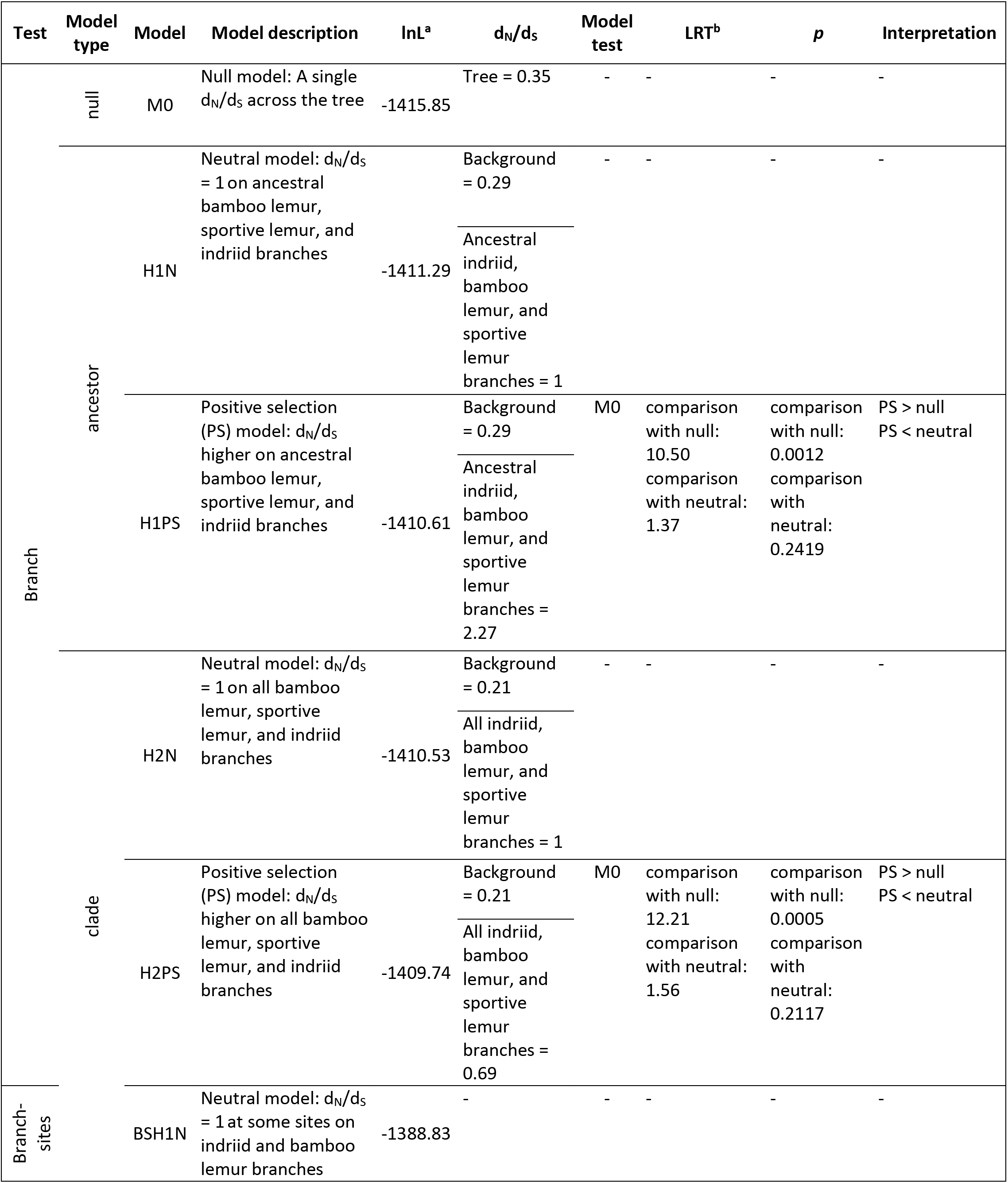
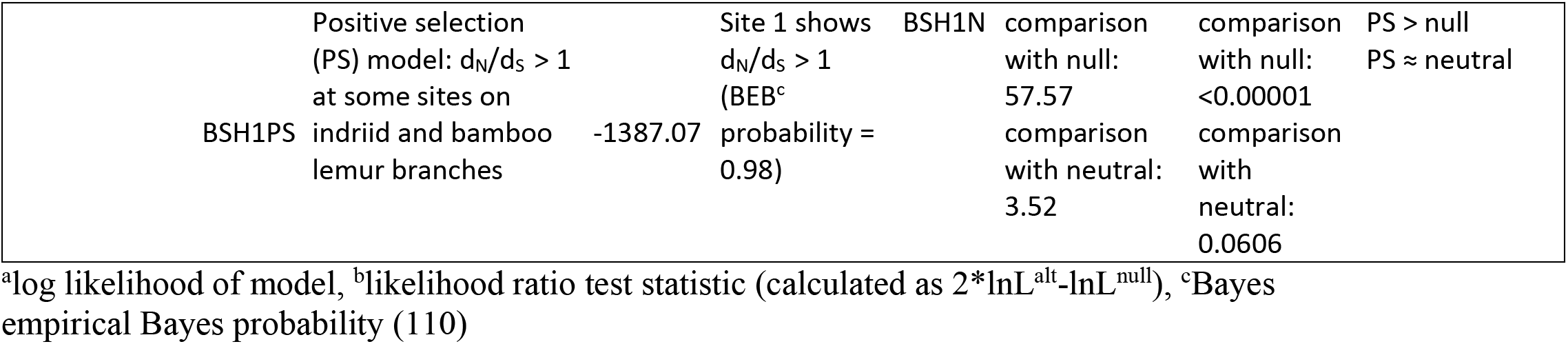
Results of codeml analysis

We include two representatives of the bamboo lemur clade, the greater bamboo lemur (*Prolemur simus*) and the eastern lesser bamboo lemur (*Hapalemur griseus*; Figure 1; Table 1). Bamboo lemurs are members of the Lemuridae family, which shares a common ancestor with indriids at least 30 Ma (Figure 1A) (3,22). Whereas most lemurids are largely omnivorous or frugivorous, bamboo lemurs are named for their dietary focus on bamboo (28). Bamboo lemurs have a relatively simple gastrointestinal tract compared with indriids and lack an enlarged cecum (27). Eastern lesser bamboo lemurs nevertheless exhibit a very long gut passage time and fiber digestibility capacity greater than sifakas (31), though greater bamboo lemurs may have a faster gut passage rate (32). Bamboo lemurs exhibit dental traits, including premolarized canines and molarized premolars, some of which are seen in other grass-eating mammals (33,34). These features likely allow them to efficiently process bamboo, which is highly fibrous and, like other grasses, gains structural integrity from silica bodies (33,34). Bamboo lemurs are also cathemeral, which has been proposed to be an adaptation to subsistence on a low quality diet, as it allows for round-the-clock intake (35,36).

The proportion of the diet comprised of bamboo varies by species, season, and geographic locale (32,37). Members of the genus *Hapalemur* appear to exhibit a degree of dietary flexibility. For example, southern bamboo lemurs (*H. meridionalis*) and eastern lesser bamboo lemur appear to successfully exploit several different habitat types and consume invasive species, including the broad-leaved paperbark tree (*Melaleuca quinquenervia*) in the case of southern bamboo lemurs (38) and guava in the case of eastern lesser bamboo lemurs (39). *Hapalemur* spp. also seem to demonstrate considerable behavioral flexibility. For example, southern bamboo lemurs increase terrestrial foraging when preferred foods are not available (40) and the Sambirano lesser bamboo lemur (*H. occidentalis*) was observed to exploit human-altered, agricultural landscapes (41). Bamboo can be a pioneer species following anthropogenic disturbance and eastern lesser bamboo lemurs have been found to occupy such edge habitats (42). It has been proposed that members of the genus *Hapalemur* are better considered ‘‘ubiquity specialists’’ or “facultative specialists” rather than bamboo specialists (37,39).

In contrast, the greater bamboo lemur is considered an “obligate specialist” (37). Madagascar giant bamboo (*Cathariostachys madagascariensis*) comprises > 95% of its diet (32). Madagascar giant bamboo is the most cyanogenic species of bamboo in Madagascar and, given its specialization on this species, greater bamboo lemurs ingest high levels of cyanide, likely necessitating physiological specialization, though the precise mechanisms of detoxification remain unknown (37). Greater bamboo lemurs focus on Madagascar giant bamboo shoots when available during the wet season and, during the dry season, are able to exploit the culm. Greater bamboo lemurs today have a very restricted and patchy geographic distribution, but the species used to be much more widespread. The lengthening of dry seasons resulting from climate change past the greater bamboo lemurs’ capacity to subsist on bamboo culm has been proposed to explain the species’ decline (43).

We also include two sportive lemurs, the weasel sportive lemur of the eastern rainforests (*Lepilemur mustelinus*) and Anjiamangirana sportive lemur (*L. grewcockorum*) of the dry, deciduous forests of the northwest. Folivory evolved at least ~12 Ma (Figure 1) in Lepilemuridae, a family of a single extant, specious genus (*Lepilemur*) (2). The 26 currently-recognized sportive lemur species (*Lepilemur* spp.) occupy nearly all forest types in Madagascar (44). They are of medium body size (600g – 1kg) (28) and are enigmatic folivores, as they push the lower boundary of expected body size for a mammal that subsists on leaves (45). Moreover, they have an energetically expensive, leaping locomotor style and are nocturnal (16). Nocturnality is rare in folivores presumably due to lower sugar content in non-photosynthesizing leaves (16) and possible increases in energetic demands of thermoregulation, especially during the cool season in Madagascar (46). Sportive lemurs further show little selectivity in leaf choice (16,46) and appear to often eat leaves that are high in alkaloids, which may function as a form of niche partitioning with similarly-sized, nocturnal, folivorous woolly lemurs (*Avahi* spp.) (13,47). Sportive lemurs have relatively short small intestines relative to other folivorous lemurs, but greatly enlarged caeca, which may allow for the selective separation and retention of digestive material and represent convergence with arboreal marsupial consumers of mature leaves (14). Their relatively simple GI tract and fairly monotonous diet of mature leaves appears to be supported by a structurally simple gut microbiota largely comprised of taxa capable of cellulolysis (29). Sportive lemurs may partly meet their energetic needs from a low-quality diet by maintaining a markedly low resting metabolic rate, even among lemurs (48). They further have a high dry matter intake for their body size (46), suggesting that they take a high-throughput strategy to nutrient acquisition, though, to our knowledge, gut passage time remains to be studied. Sportive lemurs also show potential behavioral adaptations to low nutrient intake, including very low activity levels (49), small home ranges sizes (46,50), short travel distances (16,47), and infant parking (46). Loss of contiguous habitat may thus seriously compromise their survival (47). They also may practice caecotrophy, the reingestion of feces, as exhibited by other smaller-bodied folivores like rabbits (49,51). Caecotrophy may lead to a ~50% increase in digestive efficiency and could have co-evolved with the reduction in body size from a larger-bodied, folivorous ancestor (51).

Finally, we also added representatives of the two other indriid genera, the indri (*Indri indri*) and the eastern woolly lemur (*Avahi laniger*), as well as two additional sifaka species, silky sifakas (*P. candidus*), and Milne-Edwards’ sifakas (*P. edwardsi*), thus covering the entire phylogenetic span of the indriid clade. Sifakas incorporate more fruit in their diet than do indris, which are considered flexible young leaf specialists, or woolly lemurs, which are dedicated young leaf specialists (6,17,52–55). Woolly lemurs, like sportive lemurs, are also small-bodied for folivores and nocturnal, but appear to have adopted a different folivory strategy, relying on selection of high quality leaves and exhibiting larger home ranges (13,16,47,52). Given that enlarged caeca are found among all indriids, folivory with hindgut fermentation is likely the primitive condition (28). Indriids share a common ancestor ~22-32 Ma, a time depth similar to that of the common ancestor of colobines (3,22). We thus sought to determine the pattern of amino acid substitutions over indriid evolution, including the degree to which substitutions observed in sifakas are primitive retentions from an indriid ancestor or whether species within Indriidae show parallel substitutions, as has been observed in Asian and African colobines (21).

## MATERIALS AND METHODS

### Subjects, data mining, DNA extraction, and Sanger sequencing

Our subjects were 21 species of lemur, representing five lemur families (Table 1). For several species, data came from previously released genome assemblies (56–59). We obtained the alignment of *RNASE1* from the UCSC 30-way multi-Z alignment data and BLASTed the sequence of the closest relative against available primate genomes (e.g., the *Eulemur macaco* sequence for *Prolemur simus* and *Lemur catta*; the *Microcebus murinus* sequence for *Cheirogaleus medius*). In all cases, BLAST yielded a single best hit, which we retained. We generated new data for *Avahi laniger, Indri indri*, *Propithecus candidus*, *Propithecus edwardsi, Lepilemur grewcockorum, Lepilemur mustelinus, Eulemur fulvus, Eulemur rufifrons,* and *Hapalemur griseus* (Table 1). All new data were generated from wild individuals, with the exception of the *H. griseus* individual, who was born at the Duke Lemur Center. When possible (Table 1), multiple individuals per species were analyzed to assess whether substitutions were fixed or polymorphic in the species. *P. candidus, P. edwardsi, E. fulvus,* and *E. rufifrons* samples were collected and extracted as described in (60) and *A. laniger* and *I. indri* samples as in (29). *H. griseus* liver tissue was obtained from the DLC tissue biobank, whereas *Lepilemur grewcockorum* and *Lepilemur mustelinus* samples were field collected. DNA was extracted from samples using the DNeasy Blood and Tissue Kit (Qiagen, Hilden, Germany). DNA concentrations were quantified on Qubit fluorometer (Thermo Fisher Scientific, Waltham, MA, USA). We designed primers—5’-GTCCTGTTCCCACTGCTAGTT-3’ (RN1-2-F, forward) and 5’-CGAAATGATGAGGTGGGGGTG-3’ (RN1-2-R, reverse)—to target a 489-bp portion of DNA including the coding portion of the *RNASE1* gene.The 25 μL PCR reaction included 12.5 μL Qiagen HotStartTaq Master Mix, 2.0 μL Ambion Ultrapure non-acetylated Bovine Serum Albumin (20 mg/mL), 1.0 μL each of 10 μM forward and reverse primers and 4.0 μL of template DNA. Following an activation step at 95°C for 15 min, PCR cycling conditions (35 cycles) were: 94°C for 60 sec, 60°C for 60 sec, 72°C for 60 sec. The final extension was at 72°C for 10 min. PCR products were visualized via agarose gel electrophoresis and then enzymatically purified and sequenced at the Duke DNA Analysis Facility on an Applied Biosystems 3730 Genetic Analyzer. Chromatograms were visually inspected using FinchTV v 1.5.0 (Geospiza).

### Data analysis

We assessed the occurrence of a duplication of *RNASE1* in bamboo lemurs and indriids by BLASTing (61) the *Microcebus murinus* and *Eulemur macaco RNASE1* sequences against the *Prolemur simus* (Prosim_1.0; https://www.ncbi.nlm.nih.gov/assembly/GCA_003258685.1/) and *Propithecus coquereli* (Pcoq_1.0; https://www.ncbi.nlm.nih.gov/assembly/GCF_000956105.1) genome assemblies under both “highly similar” (megablast) and “more dissimilar” (discontiguous megablast) settings. We also designed primers based on conserved regions for amplification across species included in this analysis; as such we expected that if, in the case of a gene duplication followed by divergence, primers that successfully amplified diverged *RNASE1* sequences in folivorous species would also amplify the conserved sequence, which could be evident as apparent heterozygosity in the Sanger sequencing results.

Ancestral sequences were reconstructed and amino acid substitutions mapped to a species tree using the codeml program in the PAML package (62) and visualized using the ggtree R package (63). We estimated a maximum likelihood gene tree under a general time reversible model of nucleotide substitution using RAxML v8.2.12 with the rapid bootstrap algorithm (n = 100). However, the resulting gene tree was not consistent with generally accepted genus-level relationships and seemed unlikely to reflect incomplete lineage sorting or introgression. This lack of resolution may be due to the short length of the alignment. There is not additional sequence data for this locus for some of the species included in our analyses; we were thus unable to estimate a gene tree using, for example, upstream or/and downstream DNA data. As such, we used a species tree (Figure 2A) consistent with well-supported taxonomic relationships and robustly estimated from a large genomic dataset (3) for analyses. We placed *Propithecus verreauxi*, whose taxonomic placement is uncertain, as sister to *P. coquereli* in this analysis, as their *RNASE1* sequences are identical. The only species not included in this species tree was *Propithecus candidus*. There is little genomic data available for this taxon: we added this taxon to the tree (Figure 2A) in a way consistent with best available evidence (64). Moreover, the *Propithecus candidus* sequences was identical to that of *Propithecus diadema*, which supports this placement in this analysis. We tested for positive selection on the bamboo lemur, sportive lemur, and indriid *RNASE1* sequences using branch and branch-sites models in codeml (62). Many of the branches of interest, including the ancestral bamboo lemur branch, as well as the *Propithecus tattersalli* branch and the ancestral branch leading to *A. laniger* and the *Propithecus spp.*, did not contain any synonymous substitutions, leading to d_S_ = 0, but did contain 1-3 nonsynonymous substitutions, resulting in a nonzero d_N_. These conditions make d_N_/d_S_ estimates unreliable, but likelihood ratio tests to compare models are nevertheless valid. We fit two positive selection branch models (65): H1PS (“ancestor model”) fit an elevated d_N_/d_S_ on the ancestral bamboo lemur, sportive lemur, and indriid branches (Figure 2A) and H2PS (“clade model”) fit an elevated d_N_/d_S_ on the all bamboo lemur, sportive lemur, and indriid branches. We compared this model by likelihood ratio test to a null model (M0) fitting a single d_N_/d_S_ to the entire tree and neutral models (H1N and H2N) in which d_N_/d_S_ was fixed to 1 on the foreground branches. We next used branch-sites models to test whether there might be a class of sites on the bamboo lemur, sportive lemur, and indriid branches (Figure 2A) that exhibit a d_N_/d_S_ > 1 (66). We compared this positive selection model (BSH1PS) to a neutral model (BSH1N) in which d_N_/d_S_ for this additional class of sites on these branches is fixed to 1.

To assess the potential consequences of amino acid substitutions on enzymatic activity, we calculated the isoelectric point (pI) of RNase1 for each species using the compute ExPASy pI/Mw tool (https://web.expasy.org/compute_pi/).

**Figure 2.**
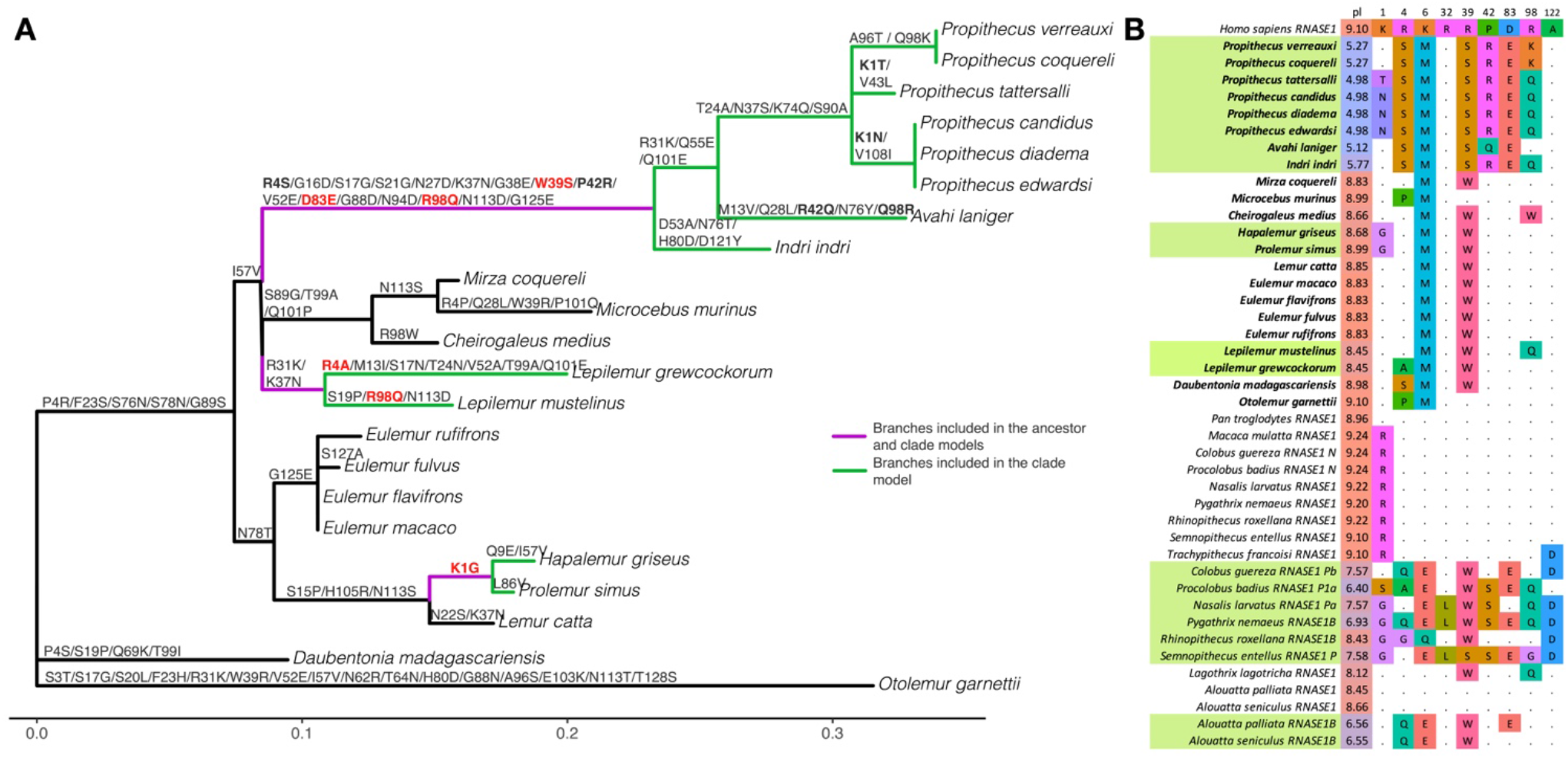
RNase1 substitutions. A = unrooted phylogenetic tree used for analyses with branches scaled to the number of substitutions per codon. Nonsynonymous substitutions mapped to tree from ancestral sequences reconstructed using codeml. Bolded substitutions occur at sites predicted to impact enzymatic activity that also show substitutions in duplicated RNase1 sequences of colobine monkeys. Substitutions in red involve changes to the same amino acid as in duplicated colobine sequences. The purple branches were the foreground branches in the ancestor positive selection branch model (H1) and branches colored both purple and green were foreground branches in the clade positive selection branch model (H2) and positive selection branch-sites model. B = Alignment of primate RNase1 sequences at sites predicted/known to influence enzymatic activity. Bolded taxon names are taxa included in the present study and taxon names highlighted in green are folivores (or the duplicated sequences of folivores in the case of colobines and howler monkeys). pI = isoelectric point at pH = 7. Primate sequences in the alignment not in the present study are from the UCSC 30-way multi-Z alignment and (19,25,67).

## RESULTS

We uncovered no evidence of a duplication of the *RNASE1* in bamboo lemurs, sportive lemurs, or indriids. We did not observe any polymorphisms within species. We did, however, observe many amino acid substitutions along the indriid branches, including at sites that influence enzymatic activity in colobine monkeys (Figure 2). In particular, these included substitutions on the ancestral indriid branch at amino acid positions 39, 83, and 98 (19,67). The substitution occurring at residue 39 involves a change from a basic to non-basic amino acid (R > S), as in colobines and howler monkeys (19,25), and is among the substitutions found to be most critical to altering enzymatic activity (67).

The indriid substitution at residue 83 involves the same exact change (D > E) as in colobines and howler monkeys (19,25) and the substitution at residue 98 involved the exact same change (R > Q) as observed in some colobines (19). We moreover observed four losses of arginines across the indriid branches; the loss of these positively-charged amino acids is thought to be especially important for RNase1 function (20).

We observed fewer amino acid substitutions on the bamboo lemur branches and no losses of arginines (Figure 2A). Further, bamboo lemurs’ RNase1 molecules do not demonstrate a lowered isoelectric point. Nevertheless, one substitution on the ancestral bamboo lemur branch occurring at residue 1 involves the same amino acid change (K > G) as observed in many duplicated colobine RNase1 sequences and that experimentally alters RNase1 enzymatic activity (19) (Figure 2).

Sportive lemurs display an intermediate scenario. They exhibit three losses of arginines, including at two sites that parallel substitutions that affect enzymatic activity in colobines (Figure 2A). One of these substitutions occurs in each of two included sportive lemur species (R4A in the Anjiamangirana sportive lemur and R98Q, also seen in indriids, in the weasel sportive lemur). An additional substitution is shared between the ancestral sportive lemur and ancestral indriid at residue 37 (K > N). The isoelectric points of sportive lemurs are slightly lower than other, non-indriid lemurs, but nevertheless much more similar to other lemurs than indriids.

The ancestor branch model (H1PS) in the codeml analysis revealed that a model including a significantly elevated d_N_/d_S_ (d_N_/d_S_ = 2.24) on the branches of the common ancestor of all bamboo lemurs, sportive lemurs, and indriids fit the data significantly better than did the null model (p = 0.0012; Table 1). However, this model was not a better fit to the data than the neutral model in which these branches have a d_N_/d_S_ = 1 (p = 0.24). The clade branch model (H2PS; Methods) including an elevated d_N_/d_S_ on all branches in the bamboo lemur, sportive lemur, and indriid clades was also better than the null model (p = 0.0005; Table 1); however, d_N_/d_S_ on these branches was not greater than 1 (d_N_/d_S_ = 0.69), indicating that most accelerated protein evolution occurred on the ancestral branches rather than in parallel within the folivore clades.

The positive selection branch-sites model identified one site (at position 1) showing a d_N_/d_S_ > 1 (Table 1). Substitutions at this site are exhibited in the bamboo lemur ancestor, the ancestor to *Propithecus candidus, P. diadema* and *P. edwardsi*, and in *P. tattersalli* (Figure 2). However, the positive selection branch-sites model was not a better fit to the data than the neutral branch-sites model (p = 0.06).

## DISCUSSION

This study reveals a probable difference in the physiological basis of folivory in the three folivorous lemur lineages, which may be relevant to better understanding the ability of these taxa to cope with habitat change. Our results indicate convergence of indriid RNase1 with duplicated RNase1s in folivorous monkeys. The elevated d_N_/d_S_ (>3) on the ancestral indriid branch together with changes in amino acids and peptide pI paralleling those in colobines and howler monkeys indicate this is a likely example of molecular adaptation in indriids. Indeed, the probability of three charge-altering amino acid substitutions occurring at the same residues in this peptide in different lineages was calculated to be extremely low (<0.0026) (21), thus emphasizing the probability of convergent adaptive function. Convergent change in *RNASE1* in indriids primarily occurred in the indriid ancestor rather than in parallel along the various indriid branches.

Evidence for adaptive convergence in the *RNASE1* gene is more limited in sportive lemurs and bamboo lemurs. Bamboo lemurs exhibit one substitution paralleling colobines and some indriids. Intriguingly, each sportive lemur lineage shows one substitution involving a loss of arginine and paralleling colobines and indriids, and the ancestral sportive lemur demonstrates additional substitutions involving losses of arginines or paralleling indriids. The results of the branch-sites tests indicate that a model including positive selection at some sites across all folivorous clades is approaching significance (*p* = 0.0606). A significantly better fit than a d_N_/d_S_ of 1 at some sites (neutral model), is a fairly conservative threshold; these results thus offer some provisional evidence for selection across all lineages.

However, neither bamboo lemurs not sportive lemurs show a substantially lowered pI. The evolutionary shift to a foliage-based diet may be somewhat more recent in sportive lemurs and is certainly more recent in bamboo lemurs than in indriids (Figure 1A). Natural selection may thus have had more time to act on *RNASE1* in indriids. *RNASE1* evolution has been explored in other mammalian bamboo specialists, specifically giant and red pandas. Notably, red pandas (*Ailurus fulgens*) exhibit a duplication of this gene with one copy characterized by numerous amino acid substitutions and a slightly lowered isoelectric point. Giant pandas (*Ailuropoda melanoleuca*), however, show no evidence of gene duplication or change in isoelectric point (68).

Alternatively, the lack of convergence among lemur lineages may reflect differing pathways to folivory. It is also noteworthy that many indriids are arguably more ecologically similar to howler monkeys and colobines than the more specialized sportive and bamboo lemurs are. Like these folivorous monkeys, indris and sifakas are fairly large-bodied and, in sifakas, in particular, flexibly incorporate some fruit in the diet (55). Molecular adaptation of *RNASE1* may thus be part of a folivory strategy involving large body size, at least in the case of indriids and howlers, caeco-colonic fermentation, and relatively generalist feeding strategies. There is some evidence that sifakas, howler monkeys, and some colobines may be somewhat resilient to habitat disturbance due to their dietary flexibility and ability to rely on foliage as fallback foods (7,69–72).

Our results generally bolster the view that folivores do not represent an ecologically uniform guild. Folivores have generally been assumed to face less food limitations and competition (73,74), which presumably make them more tolerant of habitat disturbance and climate change. This may be particularly true in the context of changing landscapes characterized by increasing anthropogenic disturbance (75). In support of this premise, extinction risk in primates has been positively associated with trophic level and negatively correlated with percent of mature leaves in the diet (76,77).

Similarly, disturbance tolerance, is negatively correlated with degree of frugivory and body size in primates (9). Anthropogenic disturbance appears to restrict dispersal in more frugivorous macaques more so than in more folivorous langurs (78).

However, increasing evidence suggests that many folivores are often quite selective foragers (79,80) and are affected behaviorally and physiologically by nutritional limitations (11,81–84). Rather than being evenly distributed, the biomass of several folivorous primates and marsupials is predicted by indicators of leaf quality, like protein to fiber ratio, secondary compound concentrations, and soil quality (4,85–89). Nutritional studies suggest that diet quality cannot be straightforwardly predicted by dietary guild, undermining the predictive utility of broad dietary categories like folivore and frugivore (90,91). Moreover, folivores generally are very specific in their choice of leaves based on species, season, tree, location within the tree, and leaf part (10,12,92). It appears that many folivores adopt strategies of either specializing on a few species or being very selective about which specimens they consume of a range of species (10).

In particular, folivores with narrower dietary breadth may have limited resilience in the face of rapid change (47,93). Specialization is associated with rarity in primates (94). In Madagascar, The levels of extinction risk of the two bamboo lemur species included here correspond to degree of specialization, with highly specialized greater bamboo lemurs classified as Critically Endangered while the more generalized eastern lesser bamboo lemurs retain a fairly broad distribution and are classified as Vulnerable (Figure 1) (95,96). Similarly, two of the more generalist folivores, sifakas and eastern lesser bamboo lemurs, show similar densities at two sites within Ranomafana National Park that differ in disturbance levels. More specialized woolly lemurs, however, show decreased densities at the more disturbed site (97). Several fairly large-bodied primate folivores appear to have highly flexible diets and incorporate a high proportion of fruit when available (90,98,99). Notably, frugo-folivores show greater variation in habitat size and patchiness, indicating potential greater tolerance for habitat disturbance, than either frugivores or folivores (100).

Understanding the many differing ways in which individual species select, process, and digest foliage is necessary to better anticipate and address resilience to habitat change. In some cases, genetic adaptation to aspects of folivory may enhance a species’ ability to cope with shifting abundances of food items (e.g., allowing a sifaka to switch between fruit and leaves), and in some cases, genetic adaptation may constrain a species to overly rely on a limited and diminishing set of food resources.

Demonstrated chemical separation of the foods selected by sympatric folivores indicates differential detoxification capacities, as well as sensory tuning to guide discrimination behavior (13,79). As such, additional genetic analyses of various genes with roles in detoxification capacity and taste and odor perception could provide insights into differences in functional capacity that could inform conservation assessments. For example, members of the cytochrome P450 monooxygenase gene superfamily are hypothesized to evolve in response to plant secondary compound composition (101), and have shown rapid evolution in folivorous mammals, including lemurs (71,102,103). Bitter taste receptor evolution may also play a key role in fine-tuning primate plant food selection (104–106). Differences in morphology are also likely to reflect differing physiological strategies. For example, gut complexity and passage time likely coevolves with the gut microbiota. Indeed, the microbiota of the sympatric folivore lemurs studied to date are markedly distinct. Phylogenetic and functional microbiome analyses have revealed that different lemur species harbor microbial communities differentially enriched for functional pathways including cellulolysis, secondary compound degradation, and amino acid metabolism (71).

It is puzzling that we found amino acid changes in RNase1 in folivorous lemurs that parallel those in duplicated RNase1 of colobines and howler monkeys without parallel gene duplications, as seen in those other primate lineages. Gene duplication can promote adaptive flexibility by allowing the original gene to retain its initial function, while duplicate copies are more likely to evolve new amino acid changes that result in novel genetic adaptation. For example, in the case of RNase1 in colobines, the enzyme encoded by the original gene is thought to play a role in the immune system by digesting double-stranded viral RNA (14). The enzymes encoded by duplicated genes in colobines have evolved amino acid changes that increase efficiency of single-stranded RNA digestion, characteristic of gut microbiota, and reduce the efficiency of double-stranded (viral) RNA digestion (13, 51). The lack of duplication in lemurs suggests that RNase1 has not both maintained its original function and gained a new function in folivorous lemurs, as in colobines and howler monkeys. Rather, given the preponderance of amino acid substitutions in indriids paralleling those in colobines, the indriid enzyme is likely to efficiently degrade the gut microbiota’s single-stranded RNA but not retain much antiviral function. In the sportive and bamboo lemurs, the situation is less clear, but the enzyme might have intermediate efficiency in digesting both microbial single-stranded and viral double-stranded RNA. This raises the question of how the enzyme’s initial, antiviral function could be lost or reduced without a prohibitive fitness cost in indriids. It may be that the digestive benefit outweighs the cost or that other genes in the RNASE gene family have taken on compensatory roles. Gene duplication in this family is not limited to evolutionary recent duplications of *RNASE1*; indeed, *RNASE1* is but one member of this gene family, which contains 13 functional genes in humans (107). The RNASE gene family’s evolution in mammals is characterized by expansion via successive duplications of an original gene and its duplicates (107). Further study is warranted to better understand the RNASE gene family evolution and enzyme function in primates.

Moreover, our results, along with the findings in howler monkeys, suggest that RNase1 plays a role in types of folivory other than strictly ruminant digestion. The proposed role of RNase1 in foregut fermentation – the reabsorption of nitrogen derived from the degradation of microbial RNA – does not explain molecular adaptation of RNase1 in hindgut fermentation. Recent work suggests that RNase1 may regulate gut microbiota composition (108). The role of RNase1 in non-ruminant folivory requires further investigation.

Ultimately, relationships between dietary guild and conservation outlook are unlikely to be straightforward. In particular, folivory probably does not indiscriminately translate into resilience to climate and habitat change. Rather, degree and specific type of specialization may be more informative. It is important to note, however, that there are likely limitations to behavioral flexibility, even for generalists or species with functional traits that “match up” with directional ecological change. For example, increased folivory may have downstream consequences for plant communities, which may result in increased defense or population crashes stemming from disruptions to pollination (37,109). Nevertheless, host genetic and microbiome studies can offer a window into physiological functional capacity that, along with behavioral, anatomical, and life history information, is relevant to conservation. This study is an example of such an approach.

## DATA AVAILABILITY STATEMENT

All accession numbers for datamined sequence data are given in Table 1. All new sequence data generated as part of this study are available on GenBank under accession numbers that are also given in Table 1.

## AUTHOR CONTRIBUTIONS

EEG and LKG designed the study, EEG, LKG, MBB, CF, and RRL conducted the genetics lab work, EEG conducted the data analysis, JR, JR, KLM, TR, HHR, REJ, CVW, ER, HAR, VR, RRL, LHA, JBC, RSR, ADY, assisted with sample collection, BJB and ADY provided reagents and supplies, EEG, LKG, MBB, RRL, and ADY wrote the manuscript, all authors read and approved the final manuscript.

## ACKNOWLEDGMENTS

We would like to thank the Marojejy guides, Ambatovy Biocamp Agents, the Bezà Mahafaly Monitoring team, Josia Razafindramanana, Vanessa Mass, Alison Richard, Patricia Wright, Thomas Gillespie, and Madagascar National Parks for facilitating sample collection in Madagascar; the Duke Lemur Center Research Department for samples; Diane Genereux and Jeremy Johnson of the Broad Institute for allowing us to use lemur genomic data they generated; members of the Duke Tung lab and Yoder lab for helpful discussion; Ziheng Yang for helpful comments on the methods; and Scott Langdon of the Duke DNA Analysis Facility for expert technical assistance. We complied with the American Association of Physical Anthropologists Code of Ethics. Our procedures were approved by the Institutional Animal Care and Use Committee of Duke University (protocols: A028-14-02; A007-17-01), and by Madagascar’s Ministère de l’Environnement, de l’Ecologie et des Forêts (MEEF/SG/DGF/DAPT/SCBT.Re permits numbers: 85/14; 68/15; 41/17; 83/17; 136/17). This is Duke Lemur Center publication #XXXXX.

